# Conceptual knowledge affects early stages of visual mental imagery and object perception

**DOI:** 10.1101/2020.01.14.905885

**Authors:** Martin Maier, Romy Frömer, Johannes Rost, Werner Sommer, Rasha Abdel Rahman

## Abstract

When we imagine an object and when we actually see that object, similar brain regions become active. Yet, the time course and mechanisms with which imagery engages perceptual networks remain to be better understood. An emerging view holds that imagery and perception follow distinct dynamics during early visual processing with similarities arising only during later, high-level visual processing. However, confounds of visual stimulation and paradigms favoring observation of high-level processes associated with subjective imagery strength may have precluded evidence of earlier shared mechanisms. We therefore manipulated prior knowledge that informs early-stage top-down predictions and tracked electrophysiological brain responses while fully controlling visual stimulation. Participants saw and imagined objects associated with varying amounts of semantic knowledge. Imagery and perception were equally influenced by knowledge at an early stage (P1 component), revealing shared mechanisms that support low-level visual processing. This finding complements previous research by showing that imagery is not merely perception in reverse. Instead, in line with the predictive processing framework, both, perception and imagery are active and constructive processes that share top-down mechanisms already in early phases of processing.

---

A substantial body of research suggests that seeing something with the mind’s eye—mental imagery—recruits similar neural circuits as seeing something with one’s physical eyes (Albers et al., 2013; Cichy et al., 2012; Dijkstra et al., 2019; Kosslyn, 2005; Kosslyn & Thompson, 2003; O’Craven & Kanwisher, 2000; Pearson, 2019; Stokes et al., 2009). This accords well with the “inside-out” view of perception proposed in the Bayesian brain-framework: In this view, perception draws on top-down predictions about incoming input that are weighed with and updated by bottom-up sensory signals. Predictions are based on a set of priors constituting an internal generative model of the external world that incorporates previous experience and knowledge (A. Clark, 2013; Friston, 2009, 2010; Hohwy, 2013; Seth, 2015). Mental imagery is initiated and sustained by top-down processing, in the absence of relevant sensory input (Dijkstra et al., 2019; Pearson, 2019). The overlap in recruited cortical networks could be explained by the notion that mental imagery largely shares the top-down aspects of perception. Differences in neural dynamics and phenomenological experience would then arise from the fact that imagery lacks the bottom-up processes propagating sensory signals that update perceptual predictions.

While overlap with perception regarding recruited cortical areas is well established, the temporal dynamics of the neurocognitive processes supporting mental imagery are less clear. According to one hypothesis, imagery may follow a reverse hierarchy compared to perception, suggesting that perception is a bottom-up process going from simple to complex visual representations, whereas imagery starts from high-level representations (e.g., of full objects) and essentially works its way down to more visual detail (Dijkstra et al., 2019; Pearson, 2019). Accordingly, during imagery, involvement of early visual cortex would become relevant only at a later stage. Supporting this view, recent studies suggested that imagery and perception show distinct neural activation patterns during early visual processing stages, which become more similar during high-level processing (Dijkstra et al., 2019; Dijkstra et al., 2018; Dijkstra, Zeidman, et al., 2017; Pearson, 2019).

However, the apparent distinct dynamics during early visual processing may result from confounds that preclude unequivocal evidence in favor of or against similarities between imagery and perception at early processing stages. First, some evidence relies on direct comparisons between imagery and perception conditions, in which visual stimulation differed substantially (e.g., Dentico et al., 2014; Dijkstra et al., 2018). Without proper controls, differences in stimulation, e.g., presenting a complex scene vs. an empty frame, trivially explain distinct neural dynamics. So far, the way to solve this visual stimulation confound has been to compare neural correlates of strength of mental imagery, for instance, how vividly a mental image appears to a participant in a given trial (Albers et al., 2013; Cui et al., 2007; Dijkstra, Bosch, et al., 2017, 2019; Fulford et al., 2018; S.-H. Lee et al., 2012). In our opinion, however, this introduces a second potential confound that specifically impairs observation of early neurocognitive mechanisms supporting imagery. The initial stages of vivid and less vivid imagery should be very similar, because in both cases participants attempt to imagine an object with the same starting conditions, e.g., in terms of initial top-down predictions. The strength of the resulting conscious mental image should depend mostly on relatively late visual processing stages that are associated with the stabilization of percepts in visual awareness and involve global recurrent processing including the frontal-parietal network (Lamme & Roelfsema, 2000; Mashour et al., 2020). These two issues can explain why imagery and perception appear to follow strictly distinct dynamics during early stages and why decodable neural patterns associated with mental imagery appear to emerge only during relatively late, high-level visual processing.

Here, we aim to complement current models by exploring early stages of mental imagery in a new way, more in line with the predictive processing view of perception and the idea that imagery essentially shares the top-down aspects of perception. The Bayesian brain-framework and previous empirical evidence contradict the notion of purely “bottom-up perception” (A. Clark, 2013; Lupyan et al., 2020; Lupyan & Clark, 2015). Perception is subject to top-down modulations already during early visual processing (Abdel Rahman & Sommer, 2008; Bar, 2004; Boutonnet & Lupyan, 2015; Collins & Olson, 2014; de Lange et al., 2018; Gandolfo & Downing, 2019; T. S. Lee & Mumford, 2003; Maier & Abdel Rahman, 2018, 2019; Press & Yon, 2019; Rabovsky et al., 2012; Samaha et al., 2018; Weller et al., 2019): For instance, semantic knowledge and conceptual-linguistic categories influence object perception as early as in the P1-component of the event-related potential (ERP), i.e. during low-level visual processing (Abdel Rahman & Sommer, 2008; Maier & Abdel Rahman, 2018, 2019; Maier et al., 2014; Rabovsky et al., 2012; Weller et al., 2019). If imagery shares the top-down aspects of perception, we hypothesize that this could hold for such early modulations as well.

Accordingly, in order to target early visual processing stages, we carefully manipulated the perceptual priors that go into imagery and perception. In a learning paradigm, we varied the depth of conceptual knowledge associated with visual objects, thus manipulating the quality of initial top-down predictions while keeping everything else constant (e.g., task, motivation, and bottom-up visual content). For the crucial comparisons, this allowed us to use the same type of stimulation (imagery or perception) and the same visual content (i.e. identical objects), and compare the influence of varying top-down predictions based on different depth of semantic object knowledge.

We combined this approach with recording and analyzing ERPs to test with high temporal precision whether early knowledge effects, repeatedly observed in perception, are mirrored in imagery. Based on previous findings (Abdel Rahman & Sommer, 2008; Rabovsky et al., 2012), we expected semantic knowledge to decrease the P1 component, a marker of sensory processing sensitive to low-level visual features such as luminance and perceived contrast (V. P. Clark et al., 1994; Di Russo et al., 2002; Foxe & Simpson, 2002; Haynes et al., 2003), as well as the N400 component, reflecting high-level semantic processing (Abdel Rahman & Sommer, 2008; Kutas & Federmeier, 2011; Rabovsky et al., 2012). We predicted that knowledge would influence both components similarly in perception and imagery.

To integrate our results with previous work, we additionally report results obtained with the established approaches of comparing imagery with perception and leveraging the strength of imagery (Dijkstra, Bosch, et al., 2017; Pearson, 2019). In line with previous evidence, we assumed that direct comparisons of perception and imagery would result in distinct dynamics during early visual processing, with increasing similarities during high-level processing. We further assumed that successful and incomplete imagery would show similar activation patterns during low-level visual processing (P1 component), but differ in high-level configural visual processing (N1 component). Combining this approach with the study of early top-down processes informed by knowledge may lead to a more complete picture of the temporal dynamics of mental imagery.

## Methods

### Participants

Participants were 32 native German speakers (23 women, 9 men; mean age 24 years; age range 20-35). All were right-handed and reported normal or corrected-to-normal visual acuity. Two participants were replaced due to excessive EEG artifacts. The study was approved by the Ethics Committee of the Psychology Department at Humboldt-Universität zu Berlin. Participants gave written informed consent and received payment or course credits.

### Apparatus and stimuli

Stimuli were presented on a 17” monitor using Presentation (Neurobehavioral Systems ®, Berkeley, USA) with a viewing distance of approximately 90 cm. The stimulus set comprised 40 rare objects (Abdel Rahman & Sommer, 2008) unfamiliar to all participants (Figure 1) and 20 well-known objects. All stimuli were gray-scale pictures of either entire objects or object fragments (used as cues), revealing about 20% of the objects, all displayed on a blue background frame of 3.5 × 3.5 cm (2.22° × 2.22° visual angle; see Figure 1). Object fragments were typical parts of the corresponding objects, allowing recognition. Fragment positions (center, left, right, top, bottom part of the object) were counterbalanced across objects. During learning, object names consisting of pseudo-nouns uninformative regarding the object’s function, were presented in both written and spoken form. In addition, for each unfamiliar object, an audio description was presented containing either a short explanation of the object’s function, use and origin (mean duration 18.3 s), or a cooking recipe (out of 20 recipes; mean duration 18.6 s, see Figure 1). Visual search displays consisted of a 7 by 7 matrix of uppercase letters with one single deviant letter. One of three different letter combinations (F-E, P-B, and T-L) was shown on a light blue background measuring 5 × 3.5 cm (3,17° × 2.22°). The deviant letter could appear in any position of the matrix except for the center column.

**Figure 1.**
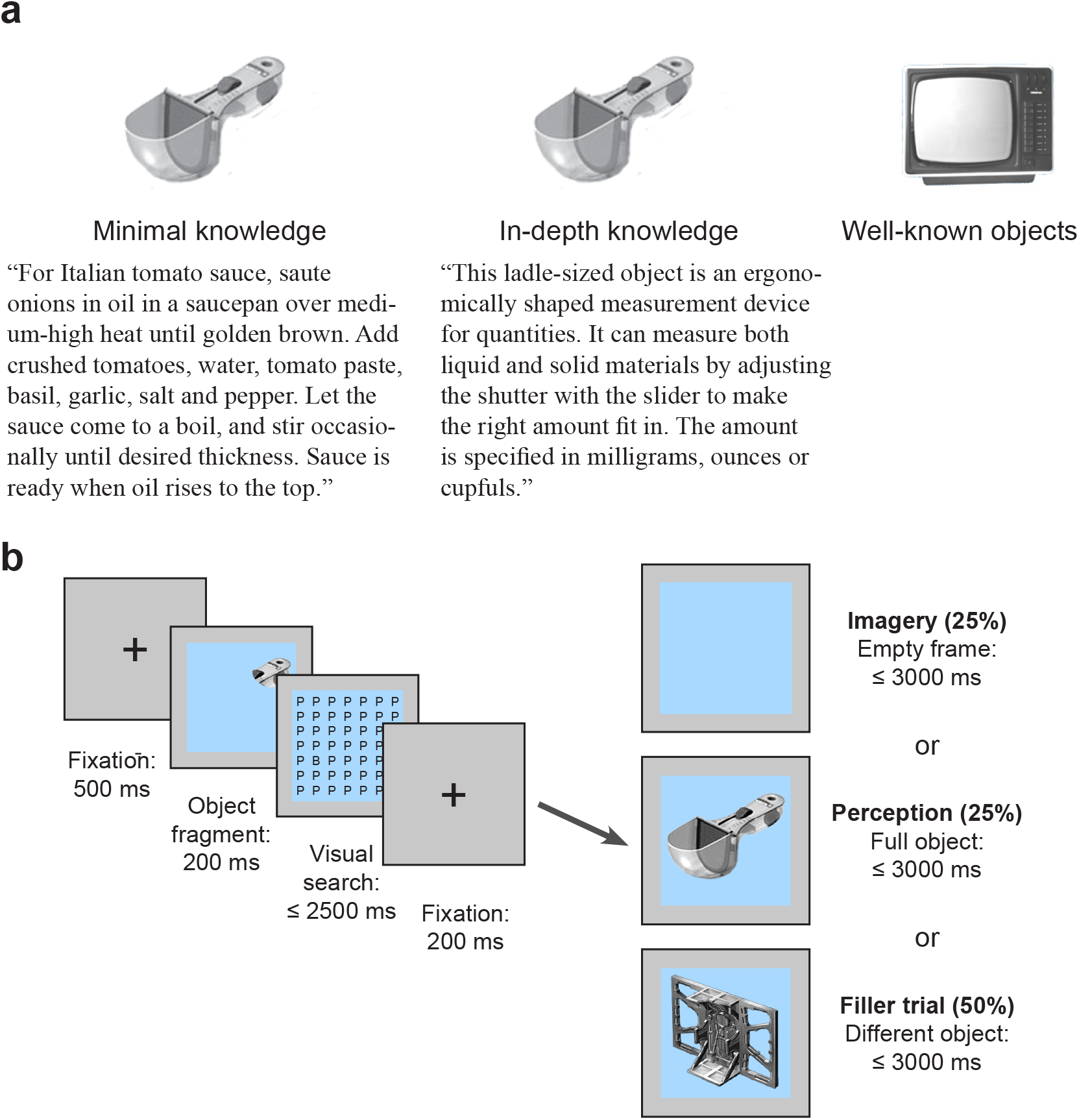
Study design. (a) Knowledge conditions with examples of object-unrelated information (minimal knowledge condition) and object-related information (in-depth knowledge condition). (b) Trial types and structure of the main task. All trial types occurred in all knowledge conditions (minimal, in-depth and well-known), with equal probability and in randomized order.

### Task and procedure

All participants completed two sessions on different days: a learning session, in which they acquired semantic knowledge about unfamiliar objects, and a test session that tested imagery and perception of the learned objects along with well-known objects.

#### Learning Phase

The learning session consisted of two parts. In Part 1, lasting about 45 minutes, participants were presented 40 unfamiliar objects and their names (written and spoken). The first part ended with a short test (approximately 10 min), comprising verbal naming and familiarity decisions on both, well-known objects and newly learned objects. In Part 2, lasting about 75 min, participants listened to recordings that provided object-related information about origin, function and use of half of the unfamiliar objects (in-depth knowledge condition), and unrelated cooking recipes for the other half (minimal knowledge condition).

Knowledge-conditions were counterbalanced across participants, such that each unfamiliar object was equally often part of the in-depth and minimal knowledge conditions.

All stories were presented twice. Thus, all unfamiliar objects were presented equally often and for the same duration and only object-related knowledge was manipulated. This resulted in three conditions with increasingly elaborate knowledge: newly learned objects without functional information (20 objects, minimal knowledge condition), newly learned objects with detailed information (20 objects, in-depth knowledge condition), and well-known objects, with preexisting information, visual and hands-on experience (20 objects, well-known objects condition). Part 2 ended with the same naming and familiarity test as Part 1.

#### Test Phase

The test session, which included EEG recordings, took place two to three days after the learning session. Before the experiment, participants filled in a knowledge questionnaire, testing recall of the pictures and related information of newly learned and well-known objects. Then, participants were familiarized with the object fragments, to make sure they could recognize the corresponding objects. Before the main task, participants performed a practice block with five well-known objects (not part of the test set), which was repeated up to two times if necessary.

In the main task, participants either imagined or saw pictures of objects. Investigating imagery with ERPs bears some timing-related difficulties: the content of imagery must be cued, but cue processing should not overlap in time with imagery, and the precise onset of imagery should be controlled. Furthermore, effects of object-knowledge on neural processing should be related to imagery, not processing of the cue.

We designed a task to control the onset and content of imagery (Figure 1). First, an object fragment was presented as cue, followed by a demanding visual search trial meant to delay the onset of imagery by taxing visual working memory (Emrich et al., 2009; Luria & Vogel, 2011; Woodman & Arita, 2011), and to avoid overlap of (semantic) processing of the cue with subsequent imagery or perception. Participants were instructed to indicate the position of a deviant letter in the left or right half of the display. Next, participants either saw an empty blue frame (imagery task, 180 trials, 25 %), a full picture of the cued object (perception task, 180 trials, 25 %), or a different object (filler trials, 360 trials, 50 %). Immediately after a response or if no response had been given within 3 s after stimulus onset, a blank screen of 1 s duration was presented. In the imagery task, participants were instructed to form an intact and detailed mental image of the cued object as quickly and accurately as possible. Participants indicated successful or incomplete imagery via button press. In perception and filler trials, participants indicated via button press whether the object was newly learned or well-known. In filler trials, two different non-corresponding object fragments were randomly assigned to each object per participant.

Requiring imagery only in 25% of the trials was aimed to discourage participants from initiating imagery already upon seeing the object fragment. Preparatory strategies were rendered ineffective by the frequent filler trials, in which invalid cues were shown. Response button assignments in the familiarity and mental imagery tasks were counterbalanced across participants. Trial types were presented in random order with short breaks after every 30 trials. Minimal knowledge, in-depth knowledge, and well-known object conditions were evenly distributed across tasks. At the end of the session, prototypical eye movements and blinks were recorded in a calibration procedure for ocular artifact correction.

### EEG recording

The EEG was recorded from 56 Ag/AgCl electrodes placed according to the extended 10-20 system, initially referenced to the left mastoid. The vertical electrooculogram (EOG) was recorded from electrodes FP1 and IO1. The horizontal EOG was recorded from electrodes F9 and F10. Electrode impedance was kept below 5 kΩ. A band pass filter with 0.032 -70 Hz, and a 50 Hz notch filter were applied; sampling rate was 250 Hz. Offline, the EEG was recalculated to average reference and low-pass filtered at 30 Hz. Eye movement and blink artifacts were removed with a spatio-temporal dipole modeling using the BESA software (Ille et al., 2002), based on the recorded prototypical eye movements and blinks. Trials with remaining artifacts and missing responses were discarded. The continuous EEG was segmented into epochs of 1.2 s locked to the stimulus of the main task (object picture or empty blue frame), including a 200 ms pre-stimulus baseline.

### Experimental Design and Statistical Analysis

Statistical analyses were performed with R (Version 4.0.0. R Core Team, 2019) and the Fieldtrip toolbox (Oostenveld et al., 2011) for Matlab (Version 2016a). Data and R-code allowing to replicate the reported analyses are available at the Open Science Framework (https://osf.io/j3vt6). Trials with unsuccessful visual search or with reaction times (RTs) shorter than 150 ms or longer than 3 SDs from individual participant’s means were excluded from all analyses. In addition, trials with incorrect familiarity classifications in the perception task were excluded from RT and ERP analyses. RTs were log transformed to approximate a normal distribution. Using the lme4 package (Version 1.1–23, Bates et al., 2015), accuracy and imagery success were analyzed with binomial generalized linear mixed models (GLMMs); RTs and ERPs were analyzed with linear mixed models (LMMs) (Frömer et al., 2018).

*P*-values were computed using the lmerTest package (Kuznetsova et al., 2016). We applied sliding difference contrasts that compare mean differences between adjacent factor levels. When indicated, we reduced models by excluding non-significant interaction terms. Model selection was performed using the anova function of the stats package in R. Along with the results of the χ^2^-Test, we compared fit indices, the Akaike information criterion (AIC) and the Bayesian information criterion (BIC), which are smaller for better model fit considering the number of parameters in each model. Behavioral data were analyzed using a nested model with the factor knowledge (well-known, in-depth and minimal) nested within task (Imagery and Perception). Mixed model analyses included random intercepts and (if supported) random slopes for subjects and object identity, allowing for better generalization of results from the particular sample of participants and the set of object pictures used here. To specify optimal random effects structures, we used the buildmer package (Voeten, 2021), which implements an algorithm identifying the maximal random effects structure with which the model still converges and performs backward stepwise elimination in order to avoid overfitting. Following Matuschek et al. (2017), both steps were performed in a backward manner (i.e. starting with the maximal model and reducing it stepwise), and likelihood ratio tests with α = .20 were conducted for model comparisons. LmerControl settings were specified to use the BOBYQA optimizer (bound optimization by quadratic approximation) with a set maximum of 200.000 iterations, and to turn off (time-consuming) derivative calculation after optimization is finished.

One follow-up analysis used Bayes factors to assess the evidence that knowledge effects on the P1 component are equal in imagery and perception. Following Vasishth et al. (2018), we used the brms package (Bürkner, 2017) to specify two different Bayesian linear mixed effects models: a null model M^0^ not assuming an interaction and an alternative model M^1^ assuming an interaction effect between task and semantic knowledge. We report the mean Bayes factor averaged from ten calculation runs, since the result can vary from run to run (Vasishth et al., 2018).

To address knowledge effects on ERPs during imagery and perception, we tested a priori hypotheses based on previous literature, that is, reduced P1 and enhanced N400 amplitudes with semantic knowledge in pre-specified regions of interest (ROIs). For the analysis of the P1 component, we averaged amplitudes within 120 to 170 ms at electrodes PO7, PO3, PO4, and PO8 (Pratt, 2011). The N400 was quantified as mean amplitude between 300 and 500 ms at electrodes PO7, PO3, PO4, PO8, O1, Oz and O2 (Abdel Rahman & Sommer, 2008; Rabovsky et al., 2012). Single trial amplitudes aggregated within ROIs and time windows were submitted to LMMs with the factors visual condition (perception, imagery and incomplete imagery) and knowledge (well-known, in-depth and minimal) as fixed effects.

To relate to previous work, we also compared imagery and perception directly and leveraged differences in the strength of imagery by comparing trials with attempted but incomplete imagery and trials with successful imagery. To this end, we calculated each participant’s average ERP in the perception, successful imagery, and incomplete imagery conditions across all scalp electrodes in time windows from 0 to 540 ms. Group-level statistics were based on paired-samples t-tests and corrected for multiple comparisons using cluster-based permutation tests (CBPT) across time and electrodes. The cluster forming threshold was set to *p* = .05. We report differences with corrected *p*-values < .025 as statistically significant.

Based on the hypothesis that imagery might be supported in particular by configural visual processing, we looked at the N1 component (Farah et al., 1988). N1 amplitudes were compared in a posterior ROI consisting of PO7, PO3, PO4, PO8, O1, Oz, and O2 (Pratt, 2011). To adjust for latency shifts (see Results), different time windows were used for the N1 component in perception and imagery, centered around the grand mean peak latencies: For perception, we aggregated over 170 – 210 ms and for imagery (both successful and incomplete) we aggregated over 210 – 250 ms. Frontal activity that coincided with the posterior N1 was analyzed in a ROI consisting of electrodes Fp1, Fpz, Fp2, AF3, AFz, AF4, F3, Fz, F4, FC1, FC2 (Gazzaley et al., 2008).

## Results

To investigate whether perception and imagery share early top-down mechanisms and to examine their time course, we recorded EEG from 32 participants while they viewed or imagined objects with varying amounts of associated knowledge. Target objects were cued with object fragments and, following an intervening visual search task to reset visual activity, participants either made a familiarity judgment about a presented object or imagined the cued object on an empty frame (see Figure 1).

### Behavioral results

In the imagery task, participants were asked to form intact and detailed mental images of the cued objects. They indicated successful or incomplete imagery via button press. Overall, participants indicated successful imagery in *M* = 84.5 % of the trials. Imagery success rates were higher in the well-known compared to the in-depth knowledge condition (*M* = 89.2 % vs. 83.0 %; nested binomial GLMM: b = 0.43, z = 3.12, *p* = .002), but there was no difference between the in-depth and the minimal knowledge condition (*M* = 83.0 % vs. 81.1 %; b = 0.08, z = 0.88, *p* = .376; see Figure 2). Knowledge affected RTs, which gradually decreased with the depth of knowledge, indicating faster imagery for objects learned with in-depth compared to minimal knowledge (*M* = 1730.4 vs. 1770.2 ms; b = -0.02, t = -2.03, *p* = .042), and for well-known objects compared to objects with in-depth knowledge (*M* = 1673.7 vs. 1730.4 ms; b = -0.05, t = -2.51, *p* = .017).

**Figure 2.**
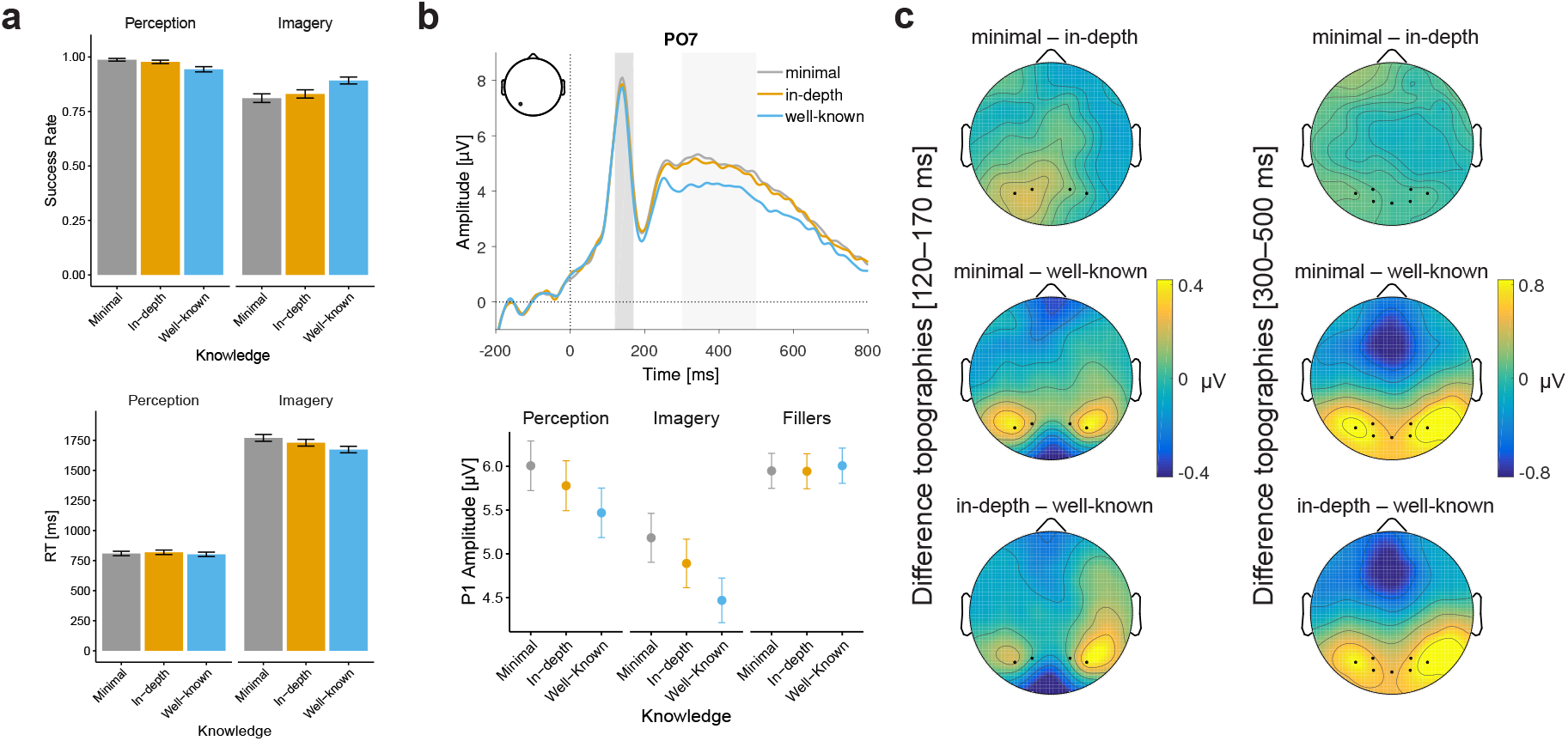
Semantic knowledge effects. (a) Behavioral results: Accuracy in the perception task and imagery success rate (top) and mean RTs (bottom) as a function of object knowledge. Error bars represent 95% confidence intervals. (b) Effects of object knowledge on the P1 and N400 components. Top panel: Grand average ERPs at electrode PO7, aggregated over perception and imagery. Bottom panel: mean P1 amplitudes by task, illustrating comparable knowledge effects in imagery and perception, and absence of the effect in filler trials. (c) Difference topographies comparing the knowledge conditions in the P1 and N400 time windows (120-170 ms; 300-500 ms, respectively). ROI electrodes are marked as dots.

In the perception task, participants classified object pictures as newly learned vs. well-known. Classification accuracy was lower in the well-known compared to the in-depth knowledge condition (*M* = 94.4 % vs. 97.8 %; b = -0.85, z = -4.02, *p* < .001) and also in the in-depth compared to the minimal knowledge condition (*M* = 97.8 % vs. 98.7 %; b = -0.55, z = -2.23, *p* = .026). RTs in the perception task (Figure 2) did not differ across knowledge conditions (in-depth -minimal: *M* = 819.0 ms vs. 810.9 ms; nested LMM: b < 0.01, t = 0.62, *p* = .534; well-known -in-depth: *M* = 801.0 ms vs. 819.0 ms; b = -0.02, t = -0.91, *p* = .371). Lower accuracy in classifying well-known objects can be explained by context effects: Participants were to classify well-known objects as “old”, but these objects had been rare during the learning session, thus in the context of the test session they were “new”. In contrast, participants were to classify newly learned objects as “new”, but in the context of the experiment these objects had been seen many times, and objects associated with richer semantic knowledge may have seemed subjectively more familiar, and thus “old”.

### Effects of semantic knowledge on ERPs

To test the hypothesis that imagery and perception share knowledge-related modulations of early visual activity, we analyzed the effects of semantic knowledge on the P1 component, an index of early perceptual processing. We further tested for later effects of knowledge in the N400, an indicator of semantic processing. In line with our hypothesis, across imagery and perception, P1 amplitudes decreased with semantic knowledge, yielding significant reductions from minimal to in-depth, and from in-depth knowledge to well-known objects (Figure 2, Table 1). The LMM including semantic knowledge, task and their interactions revealed no significant interactions of knowledge and task, suggesting similar effects of knowledge in both conditions. Exclusion of these interaction terms did not significantly decrease model fit, χ^2^(2) < 3.32, *p* > .190, and fit indices favored the reduced models (6AIC^P1^: -3, 6BIC^P1^: -18; 6AIC^N400^: 0, 6BIC^N400^: -15).

**Table 1.** Knowledge effects on the P1 and N400 components during perception and mental imagery.

| *Predictors* | **P1 component** |  |  |  | **N400 component** |  |  |  |
| --- | --- | --- | --- | --- | --- | --- | --- | --- |
|  | *Estimates* | *SE* | *t-value* | *p-value* | *Estimates* | *SE* | *t-value* | *p-value* |
| (Intercept) | 5.18 *** | 0.64 | 8.15 | <0.001 | 3.70 *** | 0.43 | 8.63 | <0.001 |
| Visual (Ima–Per) | -0.90 * | 0.38 | -2.35 | 0.024 | -0.65 | 0.35 | -1.86 | 0.072 |
| Knowledge (Deep–Min) | -0.25 * | 0.11 | -2.20 | 0.031 | -0.17 | 0.11 | -1.48 | 0.139 |
| Knowledge (Well–Deep) | -0.33 * | 0.13 | -2.56 | 0.013 | -0.46 *** | 0.13 | -3.66 | <0.001 |
| *Random Effects* | *SD* |  |  |  | *SD* |  |  |  |
| Participants | 3.41 |  |  |  | 2.38 |  |  |  |
| Visual (Ima–Per) | 2.13 |  |  |  | 1.90 |  |  |  |
| Knowledge (Deep–Min) | 0.31 |  |  |  | 0.32 |  |  |  |
| Knowledge (Well–Deep) | 0.11 |  |  |  |  |  |  |  |
| Visual × Knowledge (Deep–Min) | 0.80 |  |  |  |  |  |  |  |
| Object Identity | 0.26 |  |  |  |  |  |  |  |
| Visual (Ima–Per) | 0.66 |  |  |  |  |  |  |  |
| Residual | 4.26 |  |  |  | 4.72 |  |  |  |
| Deviance | 61256.683 |  |  |  | 63314.788 |  |  |  |
| log-Likelihood | -30628.341 |  |  |  | -31657.394 |  |  |  |
| Model | P1 ~ 1 + visual + knowledge + (1 + v_2.1 + kn_2.1 + kn_3.2 + v_2.1:kn_3.2 \| subject) + (1 + v_2.1 \| object) |  |  |  | N400 ~ 1 + visual + knowledge + (1 + v_2.1 + kn_3.2 \| subject) |  |  |  |

Note: Visual (Ima–Per) = Perception – Imagery, Knowledge (Deep–Min) = in-depth – minimal, Knowledge (Well–Deep) = well-known – in-depth
\* $p < 0.05$ \*\* $p < 0.01$ \*\*\* $p < 0.001$

Additionally, a Bayes factor analysis indicated that the observed P1-data are more likely under the assumption that knowledge effects are equal in imagery and perception. The priors going into the null model M^0^ assumed main effects of semantic knowledge (b = -0.2 for each, in-depth versus minimal knowledge and well-known vs. in-depth knowledge), but no interaction term. The alternative model M^1^ additionally assumed an interaction effect of task × semantic knowledge (b = 0.2), i.e. predicting the absence of a knowledge effect in the imagery task. The mean Bayes factor in ten calculation runs was BF^01^ = 5.85 (CI = [5.43, 6.26]), which constitutes substantial evidence in favor of M^0^, i.e. equal knowledge effects in imagery and perception.

Knowledge effects on the P1 in perception have been repeatedly observed in the absence of cueing (Abdel Rahman & Sommer, 2008; Rabovsky et al., 2012; Weller et al., 2019), and any visual priming in the present study could only occur partially as we only showed object fragments followed by an intervening visual search task to reset visual activity. Nevertheless, a potential concern with the task design is that knowledge effects on the P1 may reflect spillovers from the cues, unrelated to imagery or perception of the objects themselves. If this were true, we should observe knowledge effects also in filler trials, where different, non-cued objects were shown. In a control analysis we found no evidence that the knowledge condition of the object cue influenced the P1 in filler trials (see Figure 2 c). There was no difference between the well-known and the in-depth knowledge condition (LMM^Fillers^: b = 0.073, t = 0.600, *p* = .551) or between the in-depth and the minimal knowledge condition (LMM^Fillers^: b = -0.008, t = -0.076, *p* = .939). Thus, knowledge effects in the P1 were specific to imagining or seeing the corresponding objects.

In the N400, well-known objects produced significantly more negative amplitudes than newly learned objects, whereas the minimal and in-depth knowledge conditions did not differ (see Table 1).

To summarize, in line with our hypothesis we found semantic knowledge effects in early visual processes across both imagery and perception: P1 amplitudes were reduced with increasing depth of object-related knowledge. This effect replicates previous findings from visual perception (Abdel Rahman & Sommer, 2008; Rabovsky et al., 2012) and extends them to imagery. Previously reported differences between minimal and in-depth conditions in the N400, reflecting high-level semantic processes (Abdel Rahman & Sommer, 2008; Rabovsky et al., 2012), were not replicated.

### Comparisons between successful and incomplete imagery

To track neural activity related to imagery strength, we compared trials in which participants had indicated successful vs. incomplete imagery. The hypothesis was that incomplete compared to successful imagery may arise from failed configural processing and should thus be associated with differences in the N1 component. Since imagery may be supported by increased frontal-posterior coupling (Dentico et al., 2014; Dijkstra, Zeidman, et al., 2017), differences in frontal activity were also expected. Even though EEG scalp distributions do not translate easily to generators of activity in the brain, we hypothesized that posterior N1 effects may coincide with mirrored effects at frontal sites (Gazzaley et al., 2008). To test for global differences between successful and incomplete imagery we compared mean amplitudes with the CBPT approach, which revealed a significant difference. Underlying this difference were two clusters across electrodes and time: a posterior cluster between 228 and 392 ms, and a frontoparietal cluster between 304 and 492 ms that was slightly lateralized to the right hemisphere (Figure 3). As expected, the beginning and topography of the posterior cluster suggested a modulation of the N1 component (Figure 3).

**Figure 3.**
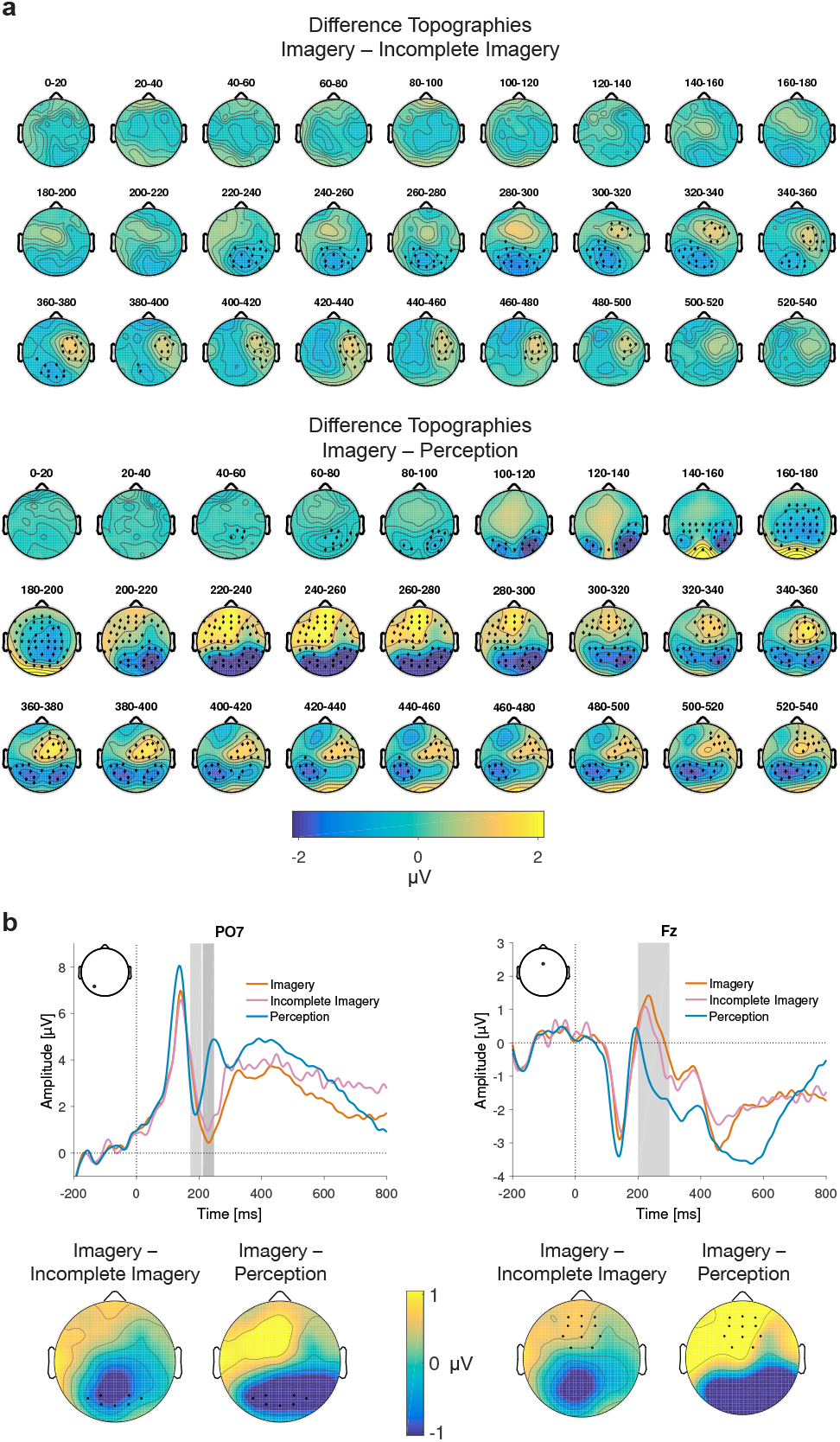
ERP comparisons of successful imagery, incomplete imagery, and perception. (a) Grand average difference-topographies. Highlighted electrodes are part of spatio-temporal clusters most compatible with the significant differences between successful and incomplete imagery (top), and between imagery and perception (bottom). (b) Comparisons of successful imagery, incomplete imagery, and perception in the N1 time window. Time windows entering the analysis of the posterior N1 amplitudes (left) and the simultaneous frontal positivity (right) are highlighted by grey shading. Topographies illustrate differences in the highlighted time windows with ROI electrodes marked by dots.

Follow-up LMM analyses based on single trial amplitudes in an independently determined posterior ROI (see Methods) confirmed a significant difference in the N1 component. Successful imagery was characterized by a larger N1 compared to incomplete imagery (Table 2). Around the same time, successful and incomplete imagery also differed at frontal sites, with a larger positivity in the frontal ROI in successful imagery trials (Figure 3, Table 2). Thus, the comparison between successful and incomplete imagery aligns with our hypothesis that successful imagery is supported by mechanisms of configural processing indexed by the posterior N1 and potentially supported by frontal top-down regulation.

**Table 2.** Comparisons between imagery, incomplete imagery, and perception in the N1 time window.

| *Predictors* | **N1 component** |  |  |  | **Frontal positivity** |  |  |  |
| --- | --- | --- | --- | --- | --- | --- | --- | --- |
|  | *Estimates* | *SE* | *t-value* | *p-value* | *Estimates* | *SE* | *t-value* | *p-value* |
| (Intercept) | 1.67 *** | 0.36 | 4.60 | <0.001 | 0.20 | 0.22 | 0.88 | 0.383 |
| Visual (Ima–Per) | -0.97 * | 0.43 | -2.24 | 0.032 | 0.62 ** | 0.21 | 2.92 | 0.006 |
| Visual (Nima–Ima) | 0.56 *** | 0.15 | 3.78 | <0.001 | -0.24 * | 0.12 | -2.04 | 0.041 |
| Centered P1 | 4.19 *** | 0.05 | 79.55 | <0.001 |  |  |  |  |
| Centered N1f |  |  |  |  | 0.71 *** | 0.01 | 81.06 | <0.001 |
| *Random Effects* | *SD* |  |  |  | *SD* |  |  |  |
| Participants | 2.03 |  |  |  | 1.22 |  |  |  |
| Visual (Ima–Per) | 2.36 |  |  |  | 1.12 |  |  |  |
| Object Identity | 0.21 |  |  |  | 0.12 |  |  |  |
| Visual (Ima–Per) | 0.58 |  |  |  | 0.31 |  |  |  |
| Visual (Nima–Ima) | 0.15 |  |  |  |  |  |  |  |
| Residual | 3.94 |  |  |  | 3.12 |  |  |  |
| Deviance | 59510.519 |  |  |  | 54515.483 |  |  |  |
| log-Likelihood | -29755.259 |  |  |  | -27257.741 |  |  |  |
| Model | N1 ~ 1 + v_2.1 + v_3.2 + cP1 + (1 + v_2.1 \| subject) + (1 + v_2.1 + v_3.2 \| object) |  |  |  | P1f ~ 1 + v_2.1 + v_3.2 + cN1f + (1 + v_2.1 \| subject) + (1 + v_2.1 \| object) |  |  |  |

Note: Visual Ima–Per = Perception – Imagery, Visual Nima–Ima = Incomplete Imagery – Imagery, P1 = preceding posterior P1 component, N1f = preceding frontal N1 component
\* $p < 0.05$ \*\* $p < 0.01$ \*\*\* $p < 0.001$

To track the neural dynamics that dissociate imagery and perception, we compared these conditions using the same approach. CBPT revealed significant differences between perception and imagery. Starting with a relative negativity for imagery at parieto-occipital sites around 80 ms post stimulus, all subsequent time windows yielded significant clusters (Figure 3). As outlined above, early differences between imagery and perception are most likely due to differences in visual stimulation. Further, differences between imagery and perception could be driven by latency shifts, amplitude differences, or both. To approximate a fair comparison, we analyzed the key ERP components—P1 and N1—in the different visual conditions at their respective peak latencies (perception, successful and incomplete imagery as one factor). P1 and N1 peak latencies were detected in the average ERP at PO7 for each participant and condition.

The LMM analysis of N1 amplitudes was adjusted for latency shifts (time windows are highlighted in Figure 3). To mitigate the influence of the different visual stimulation in imagery and perception, we included centered trial-by-trial P1 amplitudes as a covariate.

This can serve as a kind of baseline correction (Alday, 2019) because the P1 should capture a large portion of the variance related to differences in visual input and correct for amplitude differences resulting from evoked amplitude variance. The N1 was significantly larger (more negative) for successful imagery compared to perception (Table 2) and increased with more positive P1 amplitudes. Thus, the difference between perception and imagery found in the overall CBPT analysis appears to be driven by both, latency and amplitude differences.

Like for the comparison between successful and incomplete imagery, there was a modulation at frontal sites, where we found a larger positivity for imagery compared to perception coinciding with the posterior N1 component (Figure 3, Table 2). The frontal positivity further increased with more positive amplitudes of the preceding frontal negativity, which we controlled in order to account for earlier visually evoked differences.

To summarize, this second approach replicated previous work by showing distinct temporal dynamics between imagery and perception. These could largely be driven by different stimulation conditions, but latency-shifts have also added to amplitude differences. Again in line with previous studies, successful and incomplete imagery started to differ only during high-level visual processing stages. We found a larger posterior N1 for successful compared to incomplete imagery and for imagery compared to perception. These effects were accompanied by modulations of a frontal positivity in the approximate time range of the N1, which was significantly enhanced for successful compared to incomplete imagery, as well as for imagery compared to perception. Taken together these findings indicate increased demands on configural processing in imagery compared to perception, potentially supported by increased recruitment of frontal top-down processing, and that imagery fails if these increased demands are not met.

## Discussion

It is now widely accepted that visual perception and mental imagery rely on shared brain circuits, including regions in early visual cortex, as well as a network of frontal, parietal and temporal regions (Kosslyn, 2005; Pearson, 2019). Yet, the time course of imagery and the timing of the involvement of early visual cortex are still open questions. In line with predictive processing accounts we assumed that perception engages top-down predictions during low-level processing (Maier & Abdel Rahman, 2018, 2019; Maier et al., 2014; Rabovsky et al., 2012; Samaha et al., 2018; Weller et al., 2019), and hypothesized that imagery shares this mechanism.

The present work extends previous accounts holding that imagery works like perception in reverse—which assumed distinct temporal dynamics and that imagery does not rely on early perceptual representations (Dentico et al., 2014; Dijkstra et al., 2019; Dijkstra et al., 2018; Pearson, 2019). This account was supported by work showing similarities of brain activity between imagery and high-level perception (Dijkstra et al., 2018), and by imagery-related effects at the level of configural processing, as reflected in the N1 component of the ERP (Farah, 1985; Farah et al., 1988; Ganis & Schendan, 2008; Suess & Abdel Rahman, 2015). Thus, late involvement of early visual areas has been mainly supported by a lack of evidence for early involvement. Such evidence may have not been obtained, however, due to visual input confounds when comparing imagery and perception (Dijkstra et al., 2018; S.-H. Lee et al., 2012) and imagery strength manipulations favoring observation of high-level visual dynamics related to the stabilization of conscious percepts.

To overcome these obstacles, we varied the amount of knowledge associated with objects that participants saw and imagined—that is, we manipulated initial top-down predictions while controlling for bottom-up input and visual content. Using this approach, we showed that like in perception, semantic knowledge modulates low-level visual activity also during imagery, revealing similar mechanisms at a much earlier stage than previously assumed.

In line with previous evidence, we further showed that successful imagery is characterized by increased activity during high-level configural visual processing compared to both, incomplete imagery as well as perception. This suggests that demands on configural processing are higher in the absence of supporting bottom-up input. Thus, it appears that stable, high-level visual representations need to be constructed during imagery much like during perception.

### Knowledge facilitates imagery and shapes early stages of imagery and perception

In imagery, like in perception, object-related knowledge and familiarity influenced visual processing at an early stage. Deeper knowledge was associated with decreases in the amplitude of the P1 component that reflects low-level visual processing in extrastriate visual areas (V. P. Clark et al., 1994; Di Russo et al., 2002; Foxe & Simpson, 2002; Haynes et al., 2003). The influence of knowledge on the P1 in particular, with its known functional significance and localization, demonstrates that some imagery-related processes recruit early visual areas already at an early latency. Object knowledge appears to inform top-down predictions that are used in both tasks. We conclude that knowledge about an object’s function and its relevant parts facilitates low-level feature processing when we see or when we imagine that object. This novel finding suggests that imagery and perception rely on shared top-down mechanisms acting on the early construction of low-level visual representations.

Notably, the influence of semantic knowledge on imagery was of behavioral relevance: imagery of well-known objects was more often successful and faster than imagery of less familiar objects. Additionally, imagery was faster when participants had acquired in-depth rather than only minimal knowledge about initially unfamiliar objects. Thus, the more we know about an object, the better and more quickly we can imagine it.

While the P1 component in the imagery condition was evoked by a visual stimulus, the presentation of a light blue square, this physical stimulus was identical in all semantic knowledge conditions (across participants), and hence cannot have produced the observed knowledge effects. A potential objection is that modulations of early visual ERPs might not have been related to imagery, but to spillovers from the object fragment cue. However, this explanation can be ruled out because 1) the same semantic knowledge effects on perception have been shown in the absence of cueing (Abdel Rahman & Sommer, 2008; Rabovsky et al., 2012; Weller et al., 2019), 2) we only presented fragments of the objects to be imagined or perceived, 3) visual input was reset by an intervening visual search task, and, crucially, 4) there were no cue-related knowledge effects for filler trials. We therefore conclude that the influence of object knowledge on low-level visual processing is a meaningful part of imagery and perception alike.

At variance with our predictions, we did not observe an influence of in-depth versus minimal semantic knowledge on the N400 component. In contrast to previous studies demonstrating these effects in perception (Abdel Rahman & Sommer, 2008; Rabovsky et al., 2012), here, we cued the objects, which likely triggered object recognition and semantic processing. Whereas the intervening visual search task interfered with visual working memory, higher-level semantic network activation of the current object might have been sustained, given that it was potentially relevant for the upcoming task. Since the N400 is typically smaller for expected stimuli and reflects changes in semantic network activation (Rabovsky et al., 2018), the cues in our paradigm may have muted potential N400 effects.

### What distinguishes successful imagery from incomplete imagery and from perception?

In line with the notion that imagery relies strongly on high-level configural visual processing, successful and incomplete imagery started to diverge in the posterior N1 component (Busey & Vanderkolk, 2005; Costa et al., 2018; Milivojevic et al., 2008; Rossion & Jacques, 2011; Tanaka & Curran, 2001). Successful imagery was associated with larger posterior N1 amplitudes, accompanied by modulations of frontal activity at the same latency. The former finding is consistent with previous EEG and MEG studies that showed imagery-related modulations of the posterior N1 (Farah, 1985; Farah et al., 1988; Ganis & Schendan, 2008; Suess & Abdel Rahman, 2015). In terms of its functional relevance and typical latency, the N1 effect fits well with the finding that neural representations decoded from imagery using MEG match those observed in perception around 160 ms, that is, the N1 time window (Dijkstra et al., 2018). As incomplete imagery did not differ from successful imagery in the P1, it seems to share the early low-level activations but to lack (some of) the later configural processes and top-down feedback that stabilizes the percept. The reduced frontal activity associated with incomplete imagery may thus reflect insufficient involvement of frontal areas, and their connectivity to occipitotemporal visual areas, which provide crucial top-down monitoring for imagery to be maintained (Dentico et al., 2014; Dijkstra, Zeidman, et al., 2017; Pearson, 2019). Holding intact and detailed images before the mind’s eye thus seems to be supported by configural visual processing and large-scale connectivity including frontal and occipital areas that stabilizes and maintains visual representations (Dentico et al., 2014; Dijkstra et al., 2019; Dijkstra, Zeidman, et al., 2017).

This interpretation is further supported by our findings comparing imagery and perception. We found that the posterior N1 was both delayed and increased in imagery compared to perception. Simultaneously, frontal activity was more pronounced in imagery compared to perception. These results suggest that imagery relies more strongly on configural processing than perception, and engages more top-down control. When these additional demands are not met, imagery fails. To test whether success vs failure of imagery is all or none or reflects gradual degradation in configural processing, future studies could employ trial-by-trial vividness ratings to test whether they correspond to linear decreases in frontal and posterior activity (Dijkstra, Bosch, et al., 2017).

Interestingly, the finding that successful and incomplete imagery start to diverge only in the N1 component bears similarities to observations concerning conscious access and the visual awareness negativity (Förster et al., 2020). The preceding P1 component is not typically associated with perceptual awareness (Förster et al., 2020; Maier & Abdel Rahman, 2018; Sergent et al., 2005), and also did not dissociate between successful and incomplete imagery in the present study. Conscious perception is thought to depend on “global ignition” or recurrent processing in a widespread network of brain areas (Dehaene & Changeux, 2011; Lamme, 2018). It is therefore conceivable that differences between successful and incomplete imagery starting in the N1, as well as late, high-level visual representations decodable around 500 ms (Dijkstra et al., 2018), reflect the beginning of conscious mental imagery, not the beginning of imagery-related processing per se. Earlier imagery-related processing stages, revealed here by knowledge effects on the P1, appear to be pre-conscious, just as in perception.

## Conclusion

Our results provide new insights into the time course of visual mental imagery by demonstrating that top-down influences modulate imagery already at an early stage of low-level visual feature processing. This extends previous assumptions that imagery and perception share neural substrates only for high-level visual processes. Instead, they already engage common neurocognitive mechanisms during early visual processing stages—consisting in top-down predictions, informed by knowledge stored in memory.

Whether in seeing or imagining objects, our brains begin to construct what we “see” with the mind’s eye with the help of what we know.

